# Lung epithelial stem cells express SARS-CoV-2 entry factors: implications for COVID-19

**DOI:** 10.1101/2020.05.23.107334

**Authors:** Anna A. Valyaeva, Anastasia A. Zharikova, Artem S. Kasianov, Yegor S. Vassetzky, Eugene V. Sheval

## Abstract

SARS-CoV-2 can infiltrate the lower respiratory tract, resulting in severe respiratory failure and a high death rate. Normally, the airway and alveolar epithelium can be rapidly reconstituted by multipotent stem cells after episodes of infection. Here, we analyzed published RNA-seq datasets and demonstrated that cells of four different lung epithelial stem cell types express SARS-CoV-2 entry factors, including *Ace2*. Thus, stem cells can be potentially infected by SARS-CoV-2, which may lead to defects in regeneration capacity partially accounting for the severity of SARS-CoV-2 infection and its consequences.

## Introduction

The emergence of the novel SARS coronavirus 2 (SARS-CoV-2), which causes the respiratory disease COVID-19, poses an important threat to global public health. In contrast to endemic human coronaviruses, SARS-CoV-2 can infiltrate the lower respiratory tract, resulting in severe respiratory failure and a high death rate^1,2^. Infection of epithelial cells of the two major compartments in the lungs, the conducting airways and the gas-exchanging alveoli, seems to be a main cause of COVID-19 development^3^. Both the airway and alveolar epithelia are characterized by relatively slow renewal but can be rapidly reconstituted by multipotent stem cells after episodes of infection, inflammation or injury, which are commonly observed in respiratory diseases^4–6^.

To date, several types of epithelial stem cells in the distal airways and pulmonary alveoli have been identified in mice. Basal cells in the proximal airways are the major stem cell population that self-renew and, when necessary, give rise to multiple cell types, such as secretory club cells and ciliated cells^7–10^. The pulmonary alveolar epithelium is composed of two types of differentiated epithelial cells: alveolar type 1 (AT1) cells, which mediate gas exchange, and alveolar type 2 (AT2) cells, which secrete surfactant. A subpopulation of AT2 cells serves as alveolar stem cells that can differentiate into both AT1 and AT2 cells^11,12^. In addition, lung injury activates specialized epithelial stem cells residing in the distal airways to reestablish alveolar barrier function. A number of distinct, small populations of stem cells have been reported to contribute to regeneration after lung injury: variant club/secretory cells (v-club cells), bronchioalveolar stem cells (BASCs), H2-K1^high^ club cell-like stem cells and p63^+^ basal cells^6^.

Infection of epithelial stem cells can potentially lead to defects in lung regeneration capacity. Analysis of the expression of viral entry factors helps to identify human cells that can be infected by SARS-CoV-2. Cellular entry of coronaviruses depends on binding of the viral spike (S) proteins to cellular receptors and on S protein priming by host cell proteases. It was demonstrated that angiotensin-converting enzyme 2 (ACE2) is the entry receptor for SARS-CoV-2^1,13,14^, and TMPRSS2 and FURIN are major proteases for S protein priming^13,15^. Thus, (co)expression of ACE2, TMPRSS2 and FURIN is a convenient marker of cells that can be potentially infected by SARS-CoV-2^16^. Additional proteases potentially involved in SARS-CoV-2 priming include ANPEP, used by HCoV-229E^17^, and DPP4, used by MERS-CoV^18^. However, no data have been published on this subject for SARS-CoV-2.

Expression of SARS-CoV-2 entry factors can be analyzed using publicly available RNA-seq datasets. These factors, including ACE2, are highly expressed in nasal epithelial cells, but ACE2 expression on the cells of conducting airways and lung parenchyma is substantially lower^19,20^. SARS-CoV-2 entry factors are expressed in secretory and ciliated cells of the conducting airways^16,19^. In the alveolar epithelium, ACE2 expression is found only in a small subset (1-7%) of AT2 cells^16,19,21–25^, although the severity of the disease suggests a more widespread distribution. Published data on the expression of SARS-CoV-2 entry factors in stem cells are restricted to basal cells of the airway epithelium, but infection of epithelial stem cells can lead to defects in lung regeneration.

Here, to determine whether lung stem cells can be infected by SARS-CoV-2, we analyzed the expression of SARS-CoV-2 entry factors in different epithelial stem cells using publicly available RNA-seq data. Because of the limited data on lung stem cells in humans we conducted this study on data obtained from mice.

## Results

Airway epithelial cell types include basal cells, secretory club cells and ciliated cells, as well as several rare cell types – neuroendocrine, goblet and tuft cells and ionocytes. Basal cells are a heterogeneous population of stem cells of the conducting airways that can self-renew and differentiate into both secretory and ciliated epithelial lineages.

Recent studies in which publicly available scRNA-seq data were reanalyzed to detect *ACE2*-expressing cells demonstrated that only a small subpopulation of basal cells express *ACE2* and other SARS-CoV-2 entry factors; *ACE2* expression increased upon differentiation to secretory club cells^19^. We proposed that the expression of *ACE2* and other SARS-CoV-2 entry factors might be underestimated due to the presence of dropout events, i.e., a gene is detected in one cell but is not detected in another cell, usually due to extremely low mRNA input and/or the stochastic nature of gene expression^26–29^. We therefore compared datasets obtained by Montoro and coauthors using two different methods, 3′ single-cell RNA-seq (scRNA-seq) and full-length scRNA-seq^10^. Full-length scRNA-seq allows a higher number of reads per cell for each gene than 3′ scRNA-seq (396,000 reads/cell and 23,000 reads/cell, respectively^10^), which can lead to a much more precise estimation of the number of cells expressing SARS-CoV-2 entry factors.

We analyzed the expression of SARS-CoV-2 entry factors only in basal, ciliated and club cells because the number of other cells in the full-length scRNA-seq dataset was extremely low. The proportion of cells expressing *Ace2* and the priming proteases was substantially higher in the full-length scRNA-seq data (Fig. 1a). For example, the proportions of *Ace2*^+^ basal cells were 0.60% and 9.38% in the 3′ scRNA-seq and full-length scRNA-seq datasets, respectively. This indicates that the population of epithelial cells potentially sensitive to SARS-CoV-2 infection is underestimated by 3′ scRNA-seq. The expression patterns of different genes by 3′ scRNA-seq and full-length scRNA-seq were similar for highly expressed genes (e.g., *Tmprss2*), although the detected expression levels of genes with relatively low mean expression levels were substantially higher for 3′ scRNA-seq (Fig. 1b). The proportions of cells with greater-than-zero expression of different genes were extremely variable between the two datasets, particularly for genes with low and moderate expression levels; thus the dropout effect was higher for these genes (Fig. 1c). Thus, the number of cells expressing SARS-CoV-2 entry factors obtained using 3′ scRNA-seq and similar methods used in cell atlas projects could be underestimated and should be carefully interpreted.

**Figure 1.**
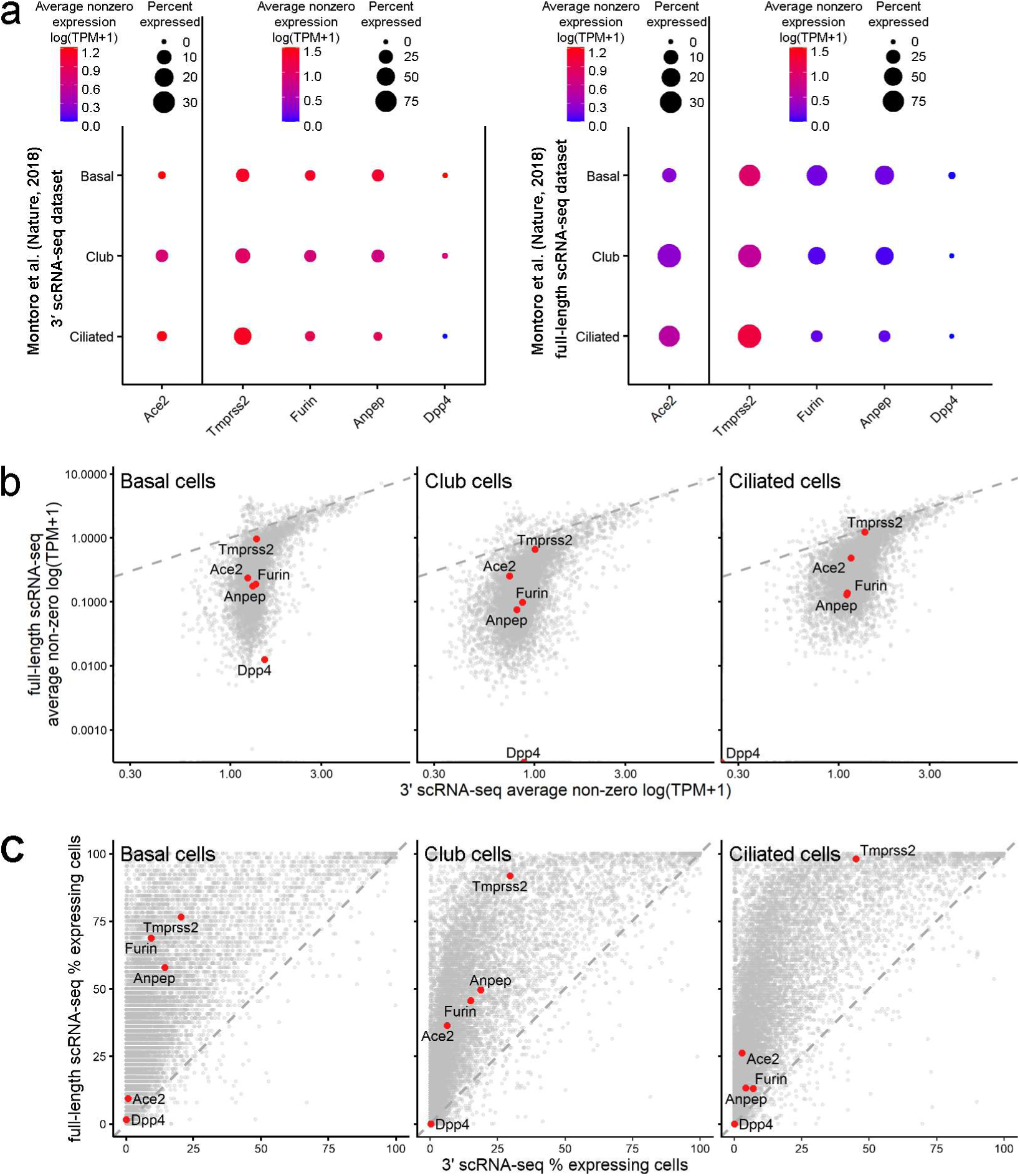
The proportions of cells expressing SARS-CoV-2 entry factors are underestimated in standard scRNA-seq datasets. (**a**) Expression of the SARS-CoV-2 entry factors *Ace2, Tmprss2, Furin, Anpep* and *Dpp4* in mouse trachea datasets from Montoro et al.^10^: 3′ scRNA-seq dataset (left panel) and full-length scRNA dataset (right panel). For the 3′ scRNA-seq dataset, unique molecular identifier (UMI) counts were normalized to account for differences in coverage, multiplied by a scaling factor of 10000 to generate transcripts per kilobase million (TPM)-like values, and then log transformed. TPM values from the full-length scRNA dataset were rescaled to sum to 10000 and were log transformed. Gene expression estimates were summarized in accordance with the cell type labels provided in the original paper. The dot size indicates the proportion of cells among the respective cell type population with greater-than-zero expression of the respective SARS-CoV-2 entry factor, while the dot color indicates the average nonzero expression value. (**b**) Correlation between gene expression in the 3′ scRNA-seq dataset and the full-length scRNA dataset for basal cells, club cells and ciliated cells. The *Ace2, Tmprss2, Furin, Anpep* and *Dpp4* expression levels are represented by colored dots. (**c**) Full-length scRNA-seq detects a substantially higher number of cells with greater-than-zero expression of genes, including SARS-CoV-2 entry factors, among basal cells, club cells and ciliated cells. The percentages of cells expressing *Ace2, Tmprss2, Furin, Anpep* and *Dpp4* are represented by the colored dots.

Only one type of lung stem cell – the basal cells of the conducting airways – has been extensively studied in both mice and humans. For the other types of stem cells, quality RNA-seq data can only be found for mice. However, the patterns of SARS-CoV-2 entry factor expression on different epithelial cells of mouse and human airways^30^ are highly similar (Supplementary Fig. 1), indicating that datasets from mice can be used for analysis of SARS-CoV-2 entry factors.

AT2 cells serve as alveolar stem cells and can differentiate into AT1 cells during alveolar homeostasis and postinjury repair^31,32^, but a small subpopulation of AT1 cells retains cellular plasticity and can transdifferentiate into AT2 cells, maintaining tissue integrity during alveolar regeneration^33,34^. These cells are characterized by the *Hopx*^+^/*Igfbp2*^*-*^ phenotype, but they do not form separated clusters in *t*-distributed stochastic neighbor embedding (*t*-SNE) plots. We analyzed the expression of SARS-CoV-2 entry factors in Igfbp2^-^ and Igfbp2^+^ AT1 cells (Fig. 2a). Neither terminally differentiated Hopx^+^/Igfbp2^+^ nor Hopx^+^/Igfbp2^-^ cells expressed *Ace2*, indicating that AT1 cells are probably resistant to SARS-CoV-2 infection. Although it is generally accepted that SARS-CoV preferentially infects AT2 cells^35^, SARS-CoV-2 can infect both AT1 and AT2 cells *ex vivo*^*36*^ and in macaques^37^. These data may indicate the presence of alternative yet unidentified pathways of SARS-CoV-2 entry into AT1 cells.

**Figure 2.**
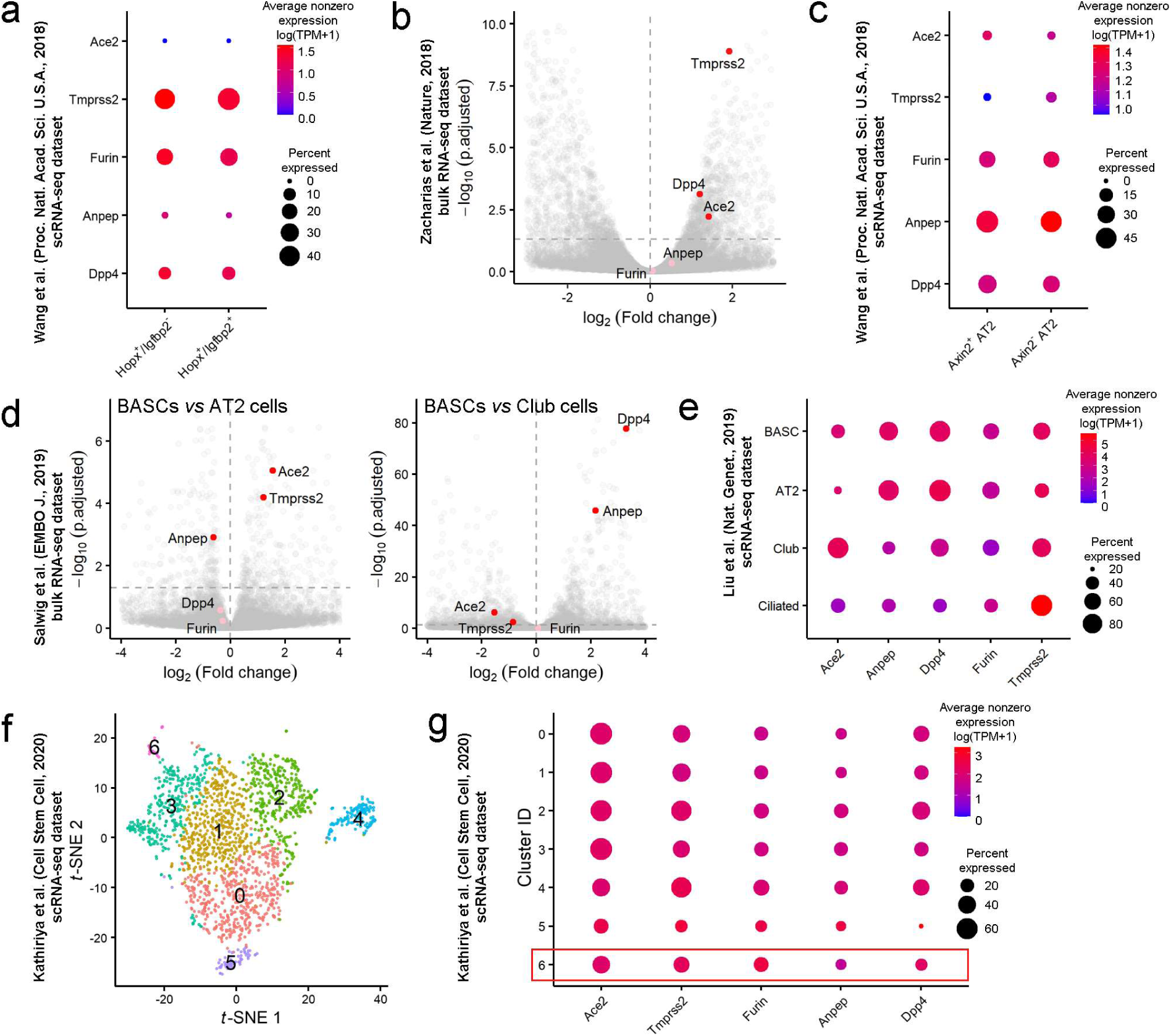
Lung epithelial stem cells express SARS-CoV-2 entry factors. (**a**) Expression of *Ace2, Tmprss2, Furin, Anpep* and *Dpp4* in the subpopulation of AT1 cells that maintain the ability to transdifferentiate into AT2 cells (*Hopx*+/*Igfbp2-*cells) and in terminally differentiated AT1 cells (*Hopx*+/*Igfbp2*+ cells). The results for the scRNA-seq dataset from mice at postnatal day 60^33^ are presented. The dot size indicates the proportion of cells in the respective cell type with greater-than-zero expression of the respective SARS-CoV-2 entry factor, while the dot color indicates the average nonzero expression value. (**b**) Volcano plot showing elevated expression of *Ace2, Tmprss2* and *Dpp4* in AEPs compared to differentiated AT2 cells (bulk RNA-seq dataset^11^). Each dot represents one gene. The log2 (fold change) in the expression levels of *Ace2, Tmprss2, Furin, Anpep* and *Dpp4* is indicated by the colored dots (red, differentially expressed genes (p.adjusted <0.05); pink, non-differentially expressed genes). (**c**) Expression of SARS-CoV-2 entry factors in AEP AT2 cells expressing *Axin2* in the dataset from Wang et al.^33^. (**d**) Volcano plots showing the expression levels of *Ace2* and *Tmprss2* in BASCs versus differentiated AT2 cells (left panel) or differentiated club cells (right panel). The log2 (fold change) in the expression levels of differentially expressed genes (p.adjusted < 0.05) is shown in red; of non-differentially expressed genes, in pink. (**e**) Expression of SARS-CoV-2 entry factors in BASCs, AT2 cells, secretory club cells and ciliated cells. The proportion of *Ace2*-expressing cells in the BASC population was higher than that in the alveolar AT2 cell population but lower than that in the bronchiolar club cell population. (**f**) t-SNE plot of 1952 scRNA-seq profiles from the scRNA-seq dataset^40^, colored by cluster assignment. Cluster 6 contains H2-K1high club-like stem cells (Supplementary Fig. 2C and 2D). (**g**) SARS-CoV-2 entry factors are expressed in a large proportion of H2-K1^high^ club-like stem cells as well as in other clusters of club cells and AT2 cells.

The major contributor to alveolar epithelial renewal is a subpopulation of AT2 cells, which serve as alveolar stem cells^11,12^. These cells, which express *Axin2*, are referred to as alveolar epithelial progenitors (AEPs)^11^. The bulk RNA-seq data for Axin2^+^ AEPs and Axin2^-^ AT2 cells^11^ were reanalyzed starting from raw reads, and transcript abundances were estimated by kallisto^38^. The expression of *Ace2, Tmprss2* and *Dpp4* was 2.68-fold, 3.79-fold and 2.31-fold, respectively, higher in AEPs than in Axin2^-^ AT2 cells (Fig. 2b). Additionally, the data on SARS-CoV-2 entry factor expression were extracted from another scRNA-seq dataset^33^. The percentage of Axin2^-^ AT2 cells with *Ace2* expression was higher than that of differentiated AT2 cells (4.20% and 2.23%, respectively), but the level of *Ace2* expression was similar in Axin2^+^ and Axin2^-^ cells (Fig. 2c). Interestingly, the number of cells expressing *Tmprss2* was low. However, both datasets indicate that alveolar stem cells (i.e., AEPs) could be even more sensitive to SARS-CoV-2 infection than differentiated AT2 cells.

The alveolar epithelium has a relatively low regeneration potential; therefore, distal airway stem cells mobilize after lung injury to occupy alveolar surfaces. Alveolar regeneration in humans after SARS-CoV-2 infection has not yet been described, but epithelial regeneration in small bronchioles has been demonstrated in a nonhuman primate model^37^. Numerous distinct populations of stem cell types have been reported to contribute to regeneration after injury^6^, but we found RNA-seq datasets for only two types of stem cells.

i. BASCs are activated and respond distinctly to different lung injuries; they also differentiate into multiple cell lineages, including club cells and ciliated cells of the terminal bronchioles and AT1 and AT2 cells of the alveoli^34,39^. Our reanalysis of a bulk RNA-seq dataset^34^ indicated that BASCs express elevated levels of *Ace2* and *Tmprss2* compared to AT2 cells (Fig. 2d, left panel). In contrast, club cells expressed higher levels of these two entry factors than BASCs (Fig. 2d, right panel). We also reanalyzed a scRNA-seq dataset^39^ to achieve better separation of the BASC cluster from other cell clusters in the *t*-SNE plot and refine the cluster labels (Supplementary Fig. 2a). BASCs coexpressed markers of AT2 cells (*Sftpc*) and club cells (*Scgb1a1*) (Supplementary Fig. 2b). Both the number of *Ace2-*expressing cells and the expression levels of *Ace2* were higher in BASCs than in AT2 cells but lower in BASCs than in club cells (Fig. 2e). Interestingly, the proportion of *Ace2*-expressing ciliated cells was roughly equal to that of BASCs, but the *Ace2* expression level in ciliated cells was substantially lower.
ii. Recently, a rare population of H2-K1^high^ club cell-like stem cells has been described. These cells, which differentiate into AT1 and AT2 cells following bleomycin-induced lung injury, were identified in scRNA-seq datasets of murine distal airways^40^. Since the cell type annotation for H2-K1^high^ cells was unavailable, we reanalyzed this dataset (Fig. 2f). Club-like cells identified by the expression of progenitor genes (Supplementary Data Fig. 2c) and secretory cell marker genes (Supplementary Fig. 2d) were reclustered into six subpopulations, including a H2-K1^high^ cell population. A relatively large proportion of H2-K1^high^ cells expressed *Ace2* (∼37.93%) and other viral entry factors, indicating that this type of stem cell is also sensitive to SARS-CoV-2 infection (Fig. 2g).

## Discussion

SARS-CoV-2 can infect lung cells and induce severe respiratory failure. Recent scRNA-seq studies demonstrated that only a minor fraction of lower respiratory tract epithelial cells expressed SARS-CoV-2 entry factors^16,19,21–25^. If the proportion of virus-sensitive cells is small, this may prevent or restrict infection of the lower respiratory tract. We suggested that the proportion of epithelial cells expressing SARS-CoV-2 entry factors could be underestimated due to the dropout effect. To roughly estimate the influence of the dropout effect, we compared two scRNA-seq datasets obtained using 3′ scRNA-seq and full-length scRNA-seq^10^ and found that the number of cells expressing *Ace2* and other SARS-CoV-2 entry factors was indeed substantially lower in the 3′ scRNA-seq dataset. For example, the proportions of *Ace2*^+^ basal cells were 0.60% and 9.38% in the 3′ scRNA-seq and full-length scRNA-seq datasets, respectively, indicating a strong influence of the dropout effect on the estimation of the number of *Ace2*-expressing cells. Hence, the fraction of cells that are potentially sensitive to the SARS-CoV-2 infection might be significantly higher than that described in the majority of published reports.

After episodes of infection or different injuries, the respiratory epithelium can be rapidly reconstituted by multipotent stem cells. Here, we analyzed published RNA-seq datasets to ascertain whether different lung epithelial stem cell types express SARS-CoV-2 entry factors. Because of the limited data on lung stem cells in humans, we conducted this study on data obtained from mice. Respiratory epithelium and epithelial stem cells are slightly different in mouse and human^6^, but the patterns of SARS-CoV-2 entry factor expression on different epithelial cells of mouse and human airways^30^ are highly similar (Supplementary Fig. 1), indicating that mouse datasets can be used for our study in the absence of valid human datasets.

We found that epithelial stem cells (basal cells, AEPs, BASCs and H2-K1^high^ cells) express *Ace*2 and other SARS-CoV-2 entry factors, making these cells probable targets of SARS-CoV-2 infection. The expression of these factors in different stem cells was relatively low, but, for example, among cells expressing markers that are specific to the gas-exchanging alveoli, AEPs exhibited higher expression of SARS-CoV-2 entry factors than differentiated AT1 and AT2 cells. These results are in agreement with the observations that cell differentiation is accompanied by depletion of the ACE2 protein^41^.

Multipotent stem cells can reconstitute lung epithelium after episodes of infection or other injuries, and it was demonstrated that stem cells could proliferate in COVID-19 patients^42^. However, the expression of SARS-CoV-2 entry factors makes them potentially infectable by SARS-CoV-2, which may in turn result in a decreased capacity for lung epithelial regeneration and potentially complicate recovery from the disease.

## Material and methods

All RNA-seq datasets used in this study were retrieved from public data repositories. Detailed information including accession codes is available in Supplementary Table 1.

### Bulk RNA-seq analysis

The data from the GSE97055 and GSE129440 datasets were reprocessed from raw reads and raw gene counts, respectively. Raw pair-end reads were quality and adapter trimmed with BBDuk (minlen=31 qtrim=r trimq=20 ktrim=r k=25 mink=11 hdist=1) from the BBTools suite, and FastQC was used for quality control. Then the reads were pseudoaligned to the mouse transcriptome (obtained from GRCm38 primary genome assembly and GENCODE gene annotation version M24 (https://www.gencodegenes.org/mouse/release_M24.html)) using kallisto^38^ with default parameters. TPM (transcripts per million) values provided by kallisto were imported and summarized into a gene-level matrix using the tximport R package^43^. Differential gene expression analysis was carried out using the DESeq2 R package^44^. Genes with Benjamini-Hochberg adjusted p-values < 0.05 were declared differentially expressed. Volcano plots were generated using the ggplot2 R package^45^.

### Single cell RNA-seq (scRNA-seq) analysis

For the GSE103354 dataset containing both 3’-droplet-based and full-length plate-based scRNA-seq data UMI (unique molecular identifier) counts and TPM values were available^10^. 3’ scRNA-seq data were normalized using the NormalizeData function from the Seurat R package^46^, and log(TP10K+1) values hereafter referred to as log(TPM+1) were obtained. TPM values of full-length scRNA-seq experiment were rescaled to add up to 10000 and log-transformed for better comparability between experiments. Further analysis of this dataset included generating average expression estimates (log of mean TPM) and percent of gene-expressing cells for clusters of basal, club, and ciliated cells based on cell labeling from the original paper^10^.

Filtered and normalized by library size (see Plasschaert et al. for details^30^) expression data from the GSE102580 dataset were log-transformed and analyzed as described above.

Raw gene counts from the GSE118891 dataset were converted to TPM values (gene lengths calculated as the union of all gene exons were obtained from Ensembl (v91) gene annotation) and log2-transformed. Cells with fewer than 2000 or more than 7500 detected genes and more than 10% of the mitochondrial fraction were excluded from the dataset. A total of 472 high-quality cells out of 480 were used for further analysis. Log2(TPM+1) values were imported into Seurat without any further normalization procedure, and a standard Seurat clustering pipeline was applied. Principal component analysis (PCA) was performed based on 2000 highly variable genes (HVGs) using the RunPCA function. The first 10 principal components (PCs) were used as the input to construct of shared nearest neighbor (SNN) graph (FindNeighbors function, k.param = 20), finding clusters (FindClusters function, resolution = 0.5) and visualizing identified clusters with the *t*-distributed stochastic neighbor embedding (*t*-SNE) algorithm (RunTSNE function, perplexity = 30). Although generated clusters were mostly consistent with cell type labels from the original paper^39^, cell type identities were refined based on the expression of known marker genes (Sftpc, Scgb1a1, Ly6a, H2-Ab, etc.) and cluster markers identified as differentially expressed genes between groups of cells by FindAllMarkers function with parameters test.use = “wilcox”, only.pos = TRUE, min.pct = 0.25, logfc.threshold = 0.25.

Log(TPM+1) expression values (scaling factor 10000) from the GSE106960 dataset were imported into Seurat with no further normalization. Since the standard cell clustering approach failed to separate the Hopx^+^/Igfbp2^-^ cell subpopulation from the bulk AT1 cells, as well as Axin2^+^ cells from AT1 cells, subclusters of interest were identified by the absolute expression of marker genes: *Hopx* and *Igfbp2* for AT1 cells and *Axin2* and *Sftpc* for AT2 cells.

For the GSE129937 dataset raw UMI counts were available. Low-quality cells were filtered out with a mitochondrial fraction threshold of 10% and a number of detected genes threshold of 1000; 3278 high-quality cells were retained for further analysis. After the data were normalized by the means of the Seurat package (NormalizeData function with default parameters), linear dimensional reduction (PCA) was conducted based on 2000 HVGs (identified by the FindVariableFeatures function with default parameters). Scores from the first 8 significant PCs (identified by inspecting the elbow plot) were used for constructing the SNN graph (FindNeighbors function, k.param = 20), finding clusters (FindClusters function, resolution = 0.4) and subsequent *t*-SNE visualization (RunTSNE function, perplexity = 50). The described analysis resulted in the identification of 10 clusters of single cells. Based on cluster marker genes obtained by the FindAllMarkers function (only.pos = TRUE, test.use = “MAST”; requires MAST R package^47^) cell type identities were assigned to clusters. Cells from 5 smaller clusters were pooled into one large cluster of secretory club-like cells based on the expression of known club cell marker genes. The pooled dataset of club-like cells were then analyzed independently. To identify the H2-K1^high^ cell population with a high progenitor gene signature supervised clustering using club cells HVGs and the list of Sox9-associated progenitor genes^48^ was carried out. A Sox9-based progenitor gene list combined with HVGs was used as a feature input for PCA. The first 10 PCs declared as significant by inspecting the elbow plot were then used to find clusters (FindNeighbors followed by FindClusters function, k.param=20, resolution = 0.5) and visualize them by the *t*-SNE algorithm (RunTSNE function, perplexity = 50). H2-K1high specific genes from Kathiriya et al.^40^ were used to locate the cluster of interest.

For all scRNA-seq datasets the average expression values (log of mean TPM) of SEFs and percent of cells expressing SEFs were calculated for every cluster of interest using the DotPlot function from Seurat.

## Code availability

R notebooks are available at https://github.com/kirushka/covid19_LSC.

## Acknowledgments

We are grateful to A.A. Penin, D.M. Potashnikova and A.A. Saidova for their helpful discussion and assistance. This work was supported by the Russian Science Foundation grant 18-14-00195 to E.V.S. computational resources of the Makarich HPC cluster were provided by the Faculty of Bioengineering and Bioinformatics, Lomonosov Moscow State University.

## Author contributions

A.A.V., A.A.Z., A.S.K. conducted the experiments, analyzed the data and wrote the paper; Y.S.V. analyzed the data and wrote the paper; E.V.S. conceived and designed the study, analyzed the data and wrote the paper.

## Supplementary Figures

**Supplementary Figure 1.**
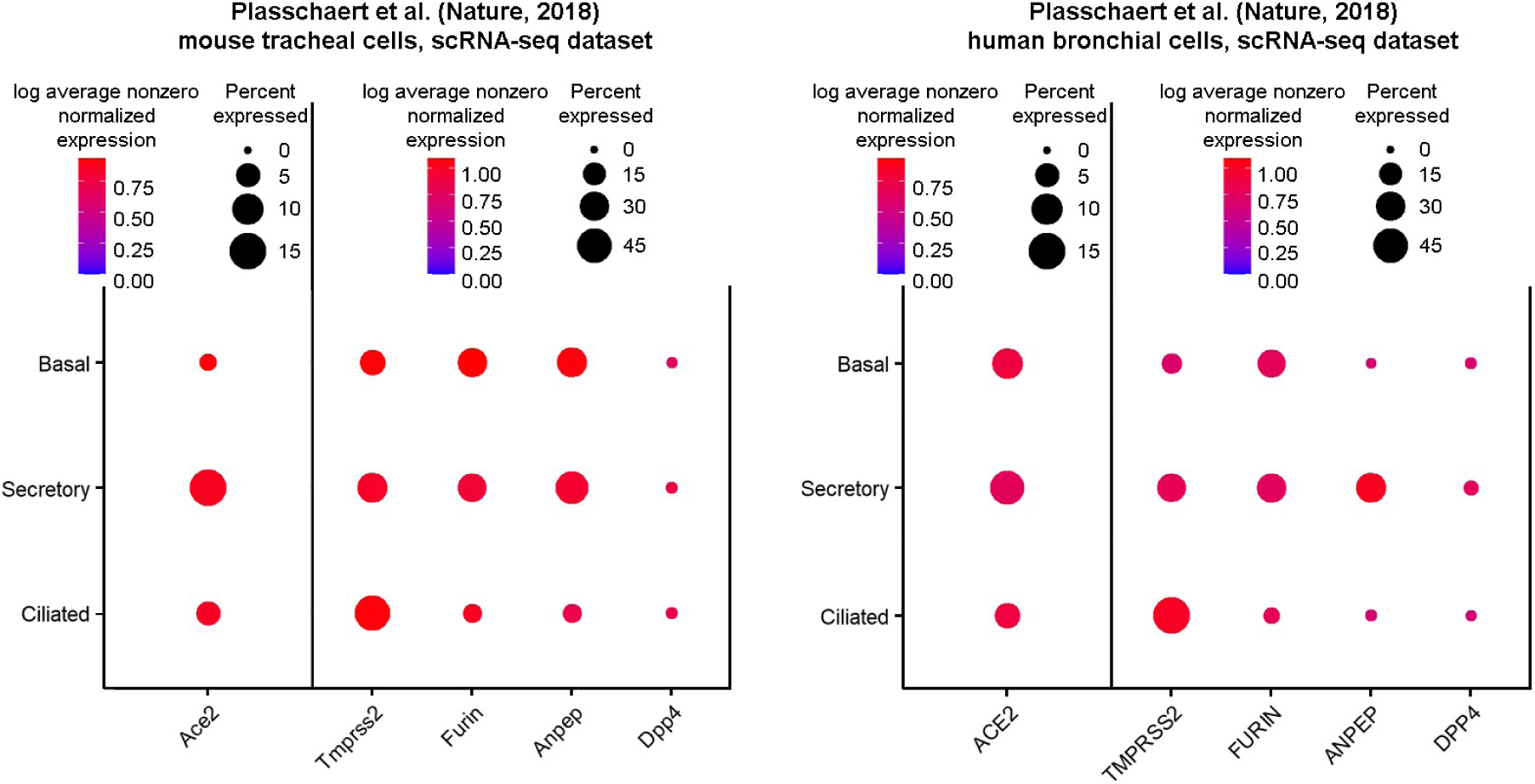
Expression of the SARS-CoV-2 entry factors in mouse and human proximal airway epithelial cells: mouse trachea scRNA-seq dataset (left panel) and human bronchiole scRNA dataset (right panel)^1^. Gene expression was estimated in accordance with the cell type labels provided in the original paper. The dot size indicates the proportion of cells among the respective cell type population with greater-than-zero expression of the respective SARS-CoV-2 entry factor, while the dot color indicates the log average nonzero expression value.

**Supplementary Figure 2.**
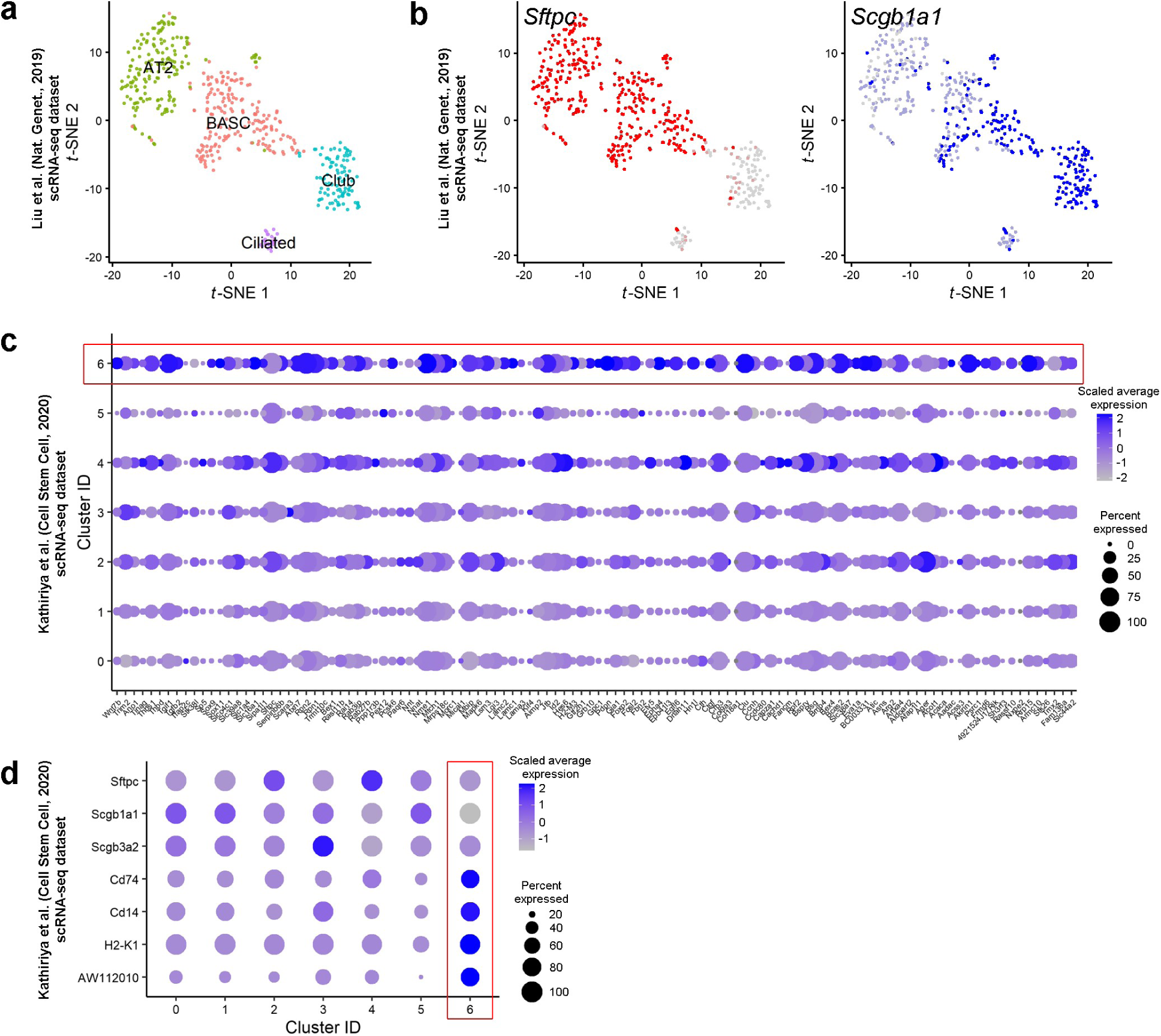
Validation of cell clusters in scRNA-seq datasets. (**a**) *t*-SNE visualization of 472 scRNA-seq profiles from the scRNA-seq dataset of FACS-sorted murine epithelial cells^2^, colored by cluster assignment and annotated *post hoc*. (**b**) *t*-SNE of 472 scRNA-seq profiles (points) colored by expression of representative AT2 cell and club cell markers (*Sftpc* and *Scgb1a1*, respectively). (**c**) Expression of progenitor cell markers^3^ in different subpopulations (1-6) of club cells^4^. Cells in Cluster 6 cells demonstrate an elevated expression of progenitor cell markers. (**d**) Expression of lineage markers of mature club cells (*Scgb1a1* and *Scgb3a2*) and AT2 cells (*Sftpc*). Note that Cluster 6 is negative or low for these markers. The cells in this cluster are characterized by enhanced expression of *Cd14, Cd74, H2-K1*, and the lncRNA AW112010s.

---

**Supplementary Table 1.** RNA-seq datasets used in this study.

| Analyzed Data | Identifier | Reference |
| --- | --- | --- |
| scRNA-seq of mouse tracheal epithelium: basal, club, ciliated, and other rare cells (7193 3' scRNA-seq cells + 301 full-length scRNA-seq cells) | GSE103354 | <sup>1</sup> |
| scRNA-seq of mouse tracheal epithelium (7661 cells from uninjured tissue) & scRNA-seq of differentiated human bronchial epithelial cells (2970 cells) | GSE102580 | <sup>2</sup> |
| bulk RNA-seq of sorted mouse AEP and AT2 cells (2 and 3 replicates per cell type) | GSE97055 | <sup>3</sup> |
| scRNA-seq of sorted mouse AT1 cells (at P3, P15, P60 stages; 3149, 2940, and 1337 cells) and AT2 cells (P60; 2093 cells) | GSE106960 | <sup>4</sup> |
| bulk RNA-seq of sorted mouse BASCs, AT2, and club cells (2 replicates per cell type) | GSE129440 | <sup>5</sup> |
| scRNA-seq of mouse BASC, AT2, and club + ciliated cells (480 cells) | GSE118891 | <sup>6</sup> |
| scRNA-seq of distal adult murine lung airway epithelium: H2-K1<sup>high</sup> cells within club-like cells (3278 cells after filtering) | GSE129937 | <sup>7</sup> |

